# Background white noise and speech facilitate visual working memory

**DOI:** 10.1101/2020.04.07.030114

**Authors:** Sizhu Han, Ruizhen Zhu, Yixuan Ku

## Abstract

In contrast to background white noise, the detrimental effects of background speech on verbal working memory (WM) were often explained by speech interference in the same verbal modality. Yet, those results were confounded with potential differences between arousal levels induced by speech and white noise. To address the role of arousal, in the present study, we minimized the verbal interference and used a visual WM task to test the influence of background speech or white noise. Electrodermal activity (EDA) and Electromyography (EMG) were recorded simultaneously to indicate the arousal levels of participants. Results showed that both background speech and white noise significantly improved visual WM performance. The change of performance further correlated with the change of physiological signals linked with arousal. Taken together, our results suggest that both background speech and white noise facilitate visual WM through raising the arousal level.

## Introduction

Playing background sound is a preferred working mode for many individuals. However, empirical studies suggest that such background sound influence human performance, either in a facilitating or a deficient way (Rauscher, Shaw, & Ky, 1993; Salamé & Baddeley, 1987; Loewen & Suedfeld, 1992; Söderlund, Sikström, & Smart, 2007).

One mechanism that underpins the deficient background sound effects on performance is the interference with working memory (WM) (Szalma & Hancock, 2011). In the context of verbal WM tasks, research found that background speech declined WM performance (Salamé & Baddeley, 1987), while background white noise did not change WM performance (Salamé & Baddeley, 1987, 1989). Although both background speech and background white noise were task-irrelevant stimuli, they could interact with the phonological loop, one slave system proposed to store phonological information in WM (Baddeley, 1996). A speech detector function was proposed within the phonological loop, interacting with background speech but not with white noise (Salamé & Baddeley, 1989), which lead to deficient effect of background speech and null-effect of background white noise. However, such model could not explain why in some cases background white noise improved WM performance (Baker & Holding, 1993; Borella et al., 2017; Salas, 2018).

A more flexible mechanism is that the background sound modulates the arousal level of participants, which decreases the breadth of attention (i.e. attentional narrowing) and further affects task performance (Broadbent, 1978; Easterbrook, 1959). In light of this account, the facilitation effect of white noise could be explained as it evokes relatively low but optimal levels of arousal and causes the individual to exclude task-irrelevant information; while the impairment caused by background speech could be explained as that it leads to relatively high but suboptimal levels of arousal, and then causes increased breadth of attention to also exclude task-relevant information. Nonetheless, the arousal level was rarely physiologically measured in background sound studies. Therefore, it is still unclear how the arousal level mediates the background sound effects and we address this issue using direct measurement of electrodermal activity (EDA) and electromyography (EMG) to track participants’ arousal changes.

It should also be noted that using verbal WM tasks includes confounding factors since background sound could interact with the phonological loop, but in the while it could influence the arousal level as well. Thus, in the present study we use visual WM tasks with Gabor patches, which minimize the interaction with the phonological loop and dissect the influence on the arousal level. In addition, besides the arousal change to bias attentional selection, using spatial cues in visual WM tasks is another way of directing attention. Specifically, a plethora of studies have indicated that a valid spatial retro-cue during the retention interval causes individual to focus on the task-relevant WM representation while ignoring the task-irrelevant WM representation and thus facilitate performance (Berryhill, Richmond, Shay, & Olson, 2012; Griffin & Nobre, 2003; Landman, Spekreijse, & Lamme, 2003; Makovski, 2012; Matsukura, Luck, & Vecera, 2007; Matsukura & Vecera, 2011; Pertzov, Bays, Joseph, & Husain, 2013; Williams & Woodman, 2012). The attentional cue can be delivered in two means: one is presented peripherally named as an exogenous cue; the other is presented centrally named as an endogenous cue (Posner, 1978). Yet, it is unclear whether background sound would modulate the retro-cueing effect, more specifically the two ways of spatial cues. Given the relationship between the arousal and the breadth of attention, we hypothesize that there would be an interaction between background sound and types of retro-cues. More specifically, we predict that background speech but not white noise would decrease the retro-cueing effect.

In sum, the present study implemented visual WM tasks (central, peripheral or no retro-cue) under two kinds of background sound (speech or white noise) or a quiet condition. Meanwhile, we used electrodermal activity (EDA) and electromyography (EMG) to track the changes in arousal level caused by the background speech or white noise, compared to the background quiet condition. We were about to investigate whether/how background sound influence visual WM through task-irrelevant ways of modulating arousal level, and whether background sound could influence the retro-cueing effect.

## Methods

### Participants

Twenty-seven healthy participants (18 - 23 years old) were recruited from East China Normal University (ECNU). Each had normal or corrected-to-normal vision and received ¥ 80 for participation. The experimental protocol was approved by human research ethics committee at ECNU, and all participants provided written informed consent prior to the experiment.

### Apparatus and stimuli

We programmed the experiment using Psychtoolbox implemented in MATLAB R2014b. Visual stimulus was presented on the screen with a resolution of 1024 * 768 pixels and 60Hz refresh rate.

We selected background white noise from the website: http://vdisk.weibo.com/s/d2wl3MjefQaw4, and speech noise from “30 Minutes News - 20180410”. Speech noise was spoken in Chinese, which was mother tongue of all participants. The length of these materials was edited using Adobe Audition software and restricted to 20 minutes. In addition, we set the volume of background speech and white noise to a fixed value beforehand, as previous research found it per se could modulate WM performance (Helps, Bamford, Sonuga-Barke, & Söderlund, 2014; G. B. Söderlund, Marklund, & Lacerda, 2009). To do so, we firstly used the keywords “white noise”, “speech noise”, “working memory” and “irrelevant speech effect” to search for relevant publications. Six out of 86 pieces of literature (see Table 1 for details) were finally selected for further analysis based on the following criteria: (1) including both quiet and noise conditions; (2) using VWM task in the experiment; (3) reporting the dB value of noise materials; (4) subjects were healthy adults.

**Table 1.** The volume of noise and corresponding performance in literatures.

| Reference | Volume(dB) | BQ | BW Raw | BW<br>Revised | BS Raw | BS Revised |
| --- | --- | --- | --- | --- | --- | --- |
| 1 | 47.5 | 2.912 | 4.228 | 0.452 | -- | -- |
| 2 | 55 | 0.17 | 0.81 | 3.765 | -- | -- |
| 3 | 57.5 | 0.28 | -- | -- | 0.17 | -0.393 |
| 4 | 70 | 28.2 | 29.6 | 0.050 | 29.9 | 0.060 |
| 5 | 78 | 40 | 31 | -0.225 | 37.5 | -0.063 |
| 6 | 85 | 37.3 | -- | -- | 36.8 | -0.013 |
|  | 90 | 37.3 | 39.3 | 0.054 | -- | -- |

*Note.* References from number 1 to 6 are in turn: Daud & Sudirman (2017), Borella et al. (2017), Sörqvist (2010), Van Gerven, Meijer, Vermeeren, Vuurman, & Jolles (2007), Söderlund & Sikström (2012) and Baker & Holding (1993). BQ = Background Quiet, BW = Background White Noise, BS = Background Speech Noise, the same below.

Data from each study was then transformed into the normalized value with the formula as below:

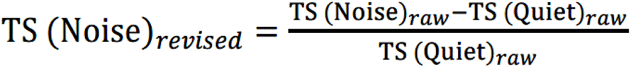

TS denotes the task score. The data after transformation (i.e., Noise revised) is shown in Table 1 and Figure 1. To maximize the reverse effects by background white noise and speech, we fixed the volume at 55dB in our experiment, measured by a decibel meter (Model AWA 5636).

**Fig. 1.**
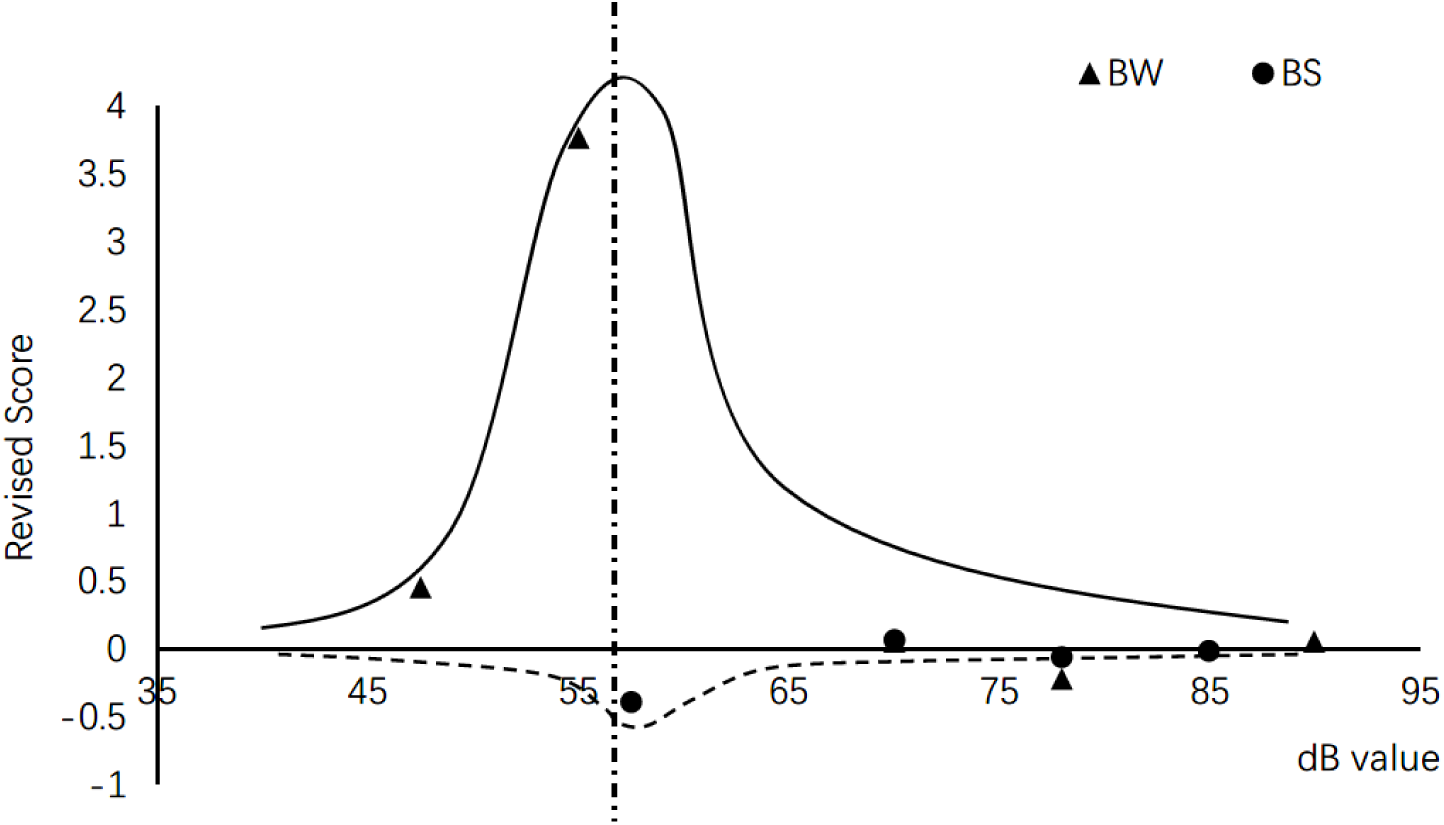
A normalized distribution of task scores as a function of volume.

### Experimental procedure

During the experiment, participants were seated 63 cm away from the screen with their head fixed on the chin rest.

As shown in Fig. 2, each trial started with a central fixation dot for 500ms, followed by memory items for another 500ms. The display consisted of two Gabor patches (radius 5°, contrast 100%, spatial frequency 2 cycles/degree) differed by at least 10°. After a short delay (2,000ms), a Gabor with a randomly chosen orientation showed up at one of the two locations where memory items were presented before. Participants were requested to adjust its orientation to match the one that was early encoded at the same location. A trial ended with a click of the left button. The inter-trial-interval was 1,000ms. In two thirds of trials, either an central (i.e., a central arrow) or an peripheral retro-cue (i.e., a peripheral circle) was presented for 100ms at 1000ms after memory items offset. In the remaining trials, the central fixation dot remained on screen during the whole delay period, without any changes to it (i.e. no cue).

**Fig. 2.**
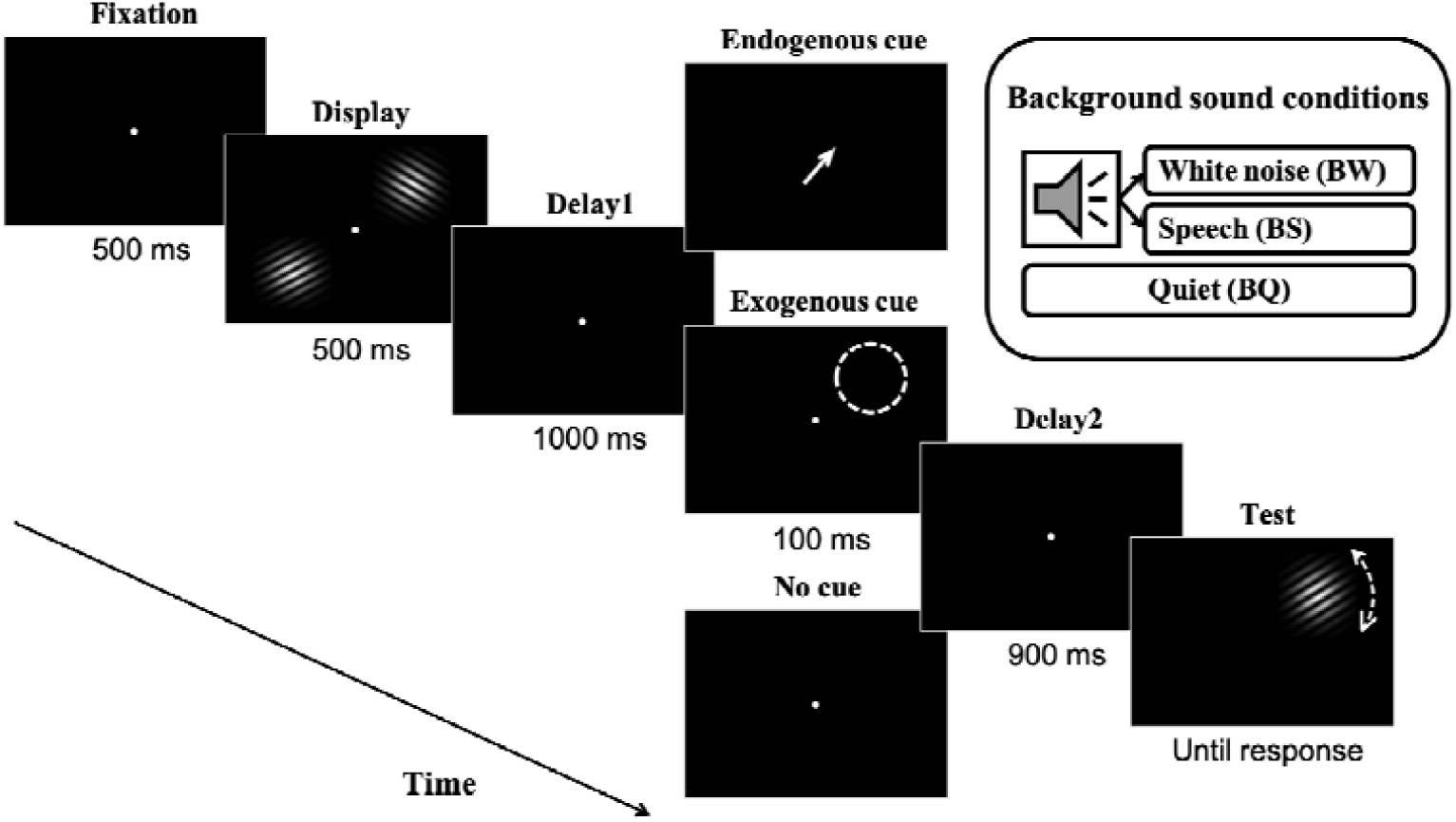
Trial sequence. After a fixation dot, two Gabor patches were presented followed by an initial delay period. Next, one of three cues (a central arrow indicates the central cue, a dashed circle at the periphery indicates the peripheral cue, a fixation point indicates the no cue) flashed and followed by a second delay period. After the second delay period, a probed Gabor appeared and subjects had to rotate the orientation of the Gabor to match the one in their memory at the same location by using the mouse.

To get familiar with the task, participants were instructed to practice 20 trials. The experiment contained 9 blocks with 72 trials each. Three cue types were randomly intermixed within each block. The auditory condition alternated in every 3 blocks, whose sequence was counterbalanced across participants. Background sound was played at the beginning of each block and stopped by the experimenter at the end of the block.

### Electrophysiological data collection

We used MP150 multi-conducting physiological recorder (America, BIOPAC company) to collect electrophysiological signals accompanied by sound. Specifically, EMG signals were measured through an EMG100C amplifier with 2 shielded LEAD11A0S wires, 1 non-shielded LEAD100A wire and 3 BIOPAC disposable electrodes. EDA signals were measured through a skin electric sensor (TSD203). These signals for each block were separately recorded on AcqKnowledge 5.0 software and then exported for further analysis.

## Data analysis

### Behavioral analysis

We calculated Raw SD as the circular standard deviation of response errors (−90° ∼ 90°), which was adjusted with CircStat2012a toolbox (Berens, 2009). Two sources of response errors, the guess rate and SD (1/precision), were respectively estimated from the Standard Mixture Model (Zhang & Luck, 2008) using the MemToolbox toolbox (Suchow, Brady, Fougnie, & Alvarez, 2013). The guess rate represents the probability that the target was not in memory. The guess rate of 0.5 served as a chance level that participants make responses at random. SD indicates the precision of the target that was in memory. Three subjects were excluded due to poor behavioral performance (guess rate>0.5).

### Electrophysiological analysis

EMG and EDA reflect the electrical impulses of muscle fibers and the skin conductance, respectively. A lower value of EMG or a higher value of EDA suggest a larger level of arousal (Thompson, Mackenzie, Leuthold, & Filik, 2016). We calculated averaged EMG and EDA signals for each auditory condition. Data from 4 participants was additionally excluded due to abnormal values (>2 standard deviations) or missing data (2 subjects), leaving data from 20 subjects were included into analysis.

### Statistical analysis

All analysis was performed using the SPSS 22.0 software. For behavioral data, we performed a two-way repeated ANOVA with auditory condition and visual cue type as factors. For physiological data, we performed one-way repeated ANOVA with auditory condition as a factor. Post-hoc *t*-tests were further conducted if significant main effects or interactions were founded (alpha<0.05). Finally, we used Pearson correlation to analyze the correlation between behavioral and physiological data.

## Results

### Behavioral results

#### Raw SD

As shown in Fig. 3a, there was no interaction between auditory condition and cue type (*F* (4, 76) =0.644, *p*=0.633, η^2^=0.033). The main effect of auditory condition was significant (*F* (2, 38) =8.539, *p*=0.001, η^2^=0.310). Post hoc *t*-tests (see Fig. 3b) revealed that the Raw SD was significantly lower in background white noise (*t*=-3.092, *p*=0.006) and speech noise (*t*=-3.547, *p*=0.002) than in background quiet condition, but no difference was found between background white noise and speech noise condition (*p*=0.830). The main effect of cue type was also significant (*F* (2, 38) =16.972, *p*<0.001, η^2^ =0.472). Post hoc *t*-tests (see Fig. 3c) revealed that Raw SD was significantly lower in both peripheral (*t*=-4.547, p<0.001) and central retro-cues (*t*=-4.417, *p*<0.001) compared to no cue condition, there was no significant difference between two types of cues (*p*=0.863).

**Fig. 3.**
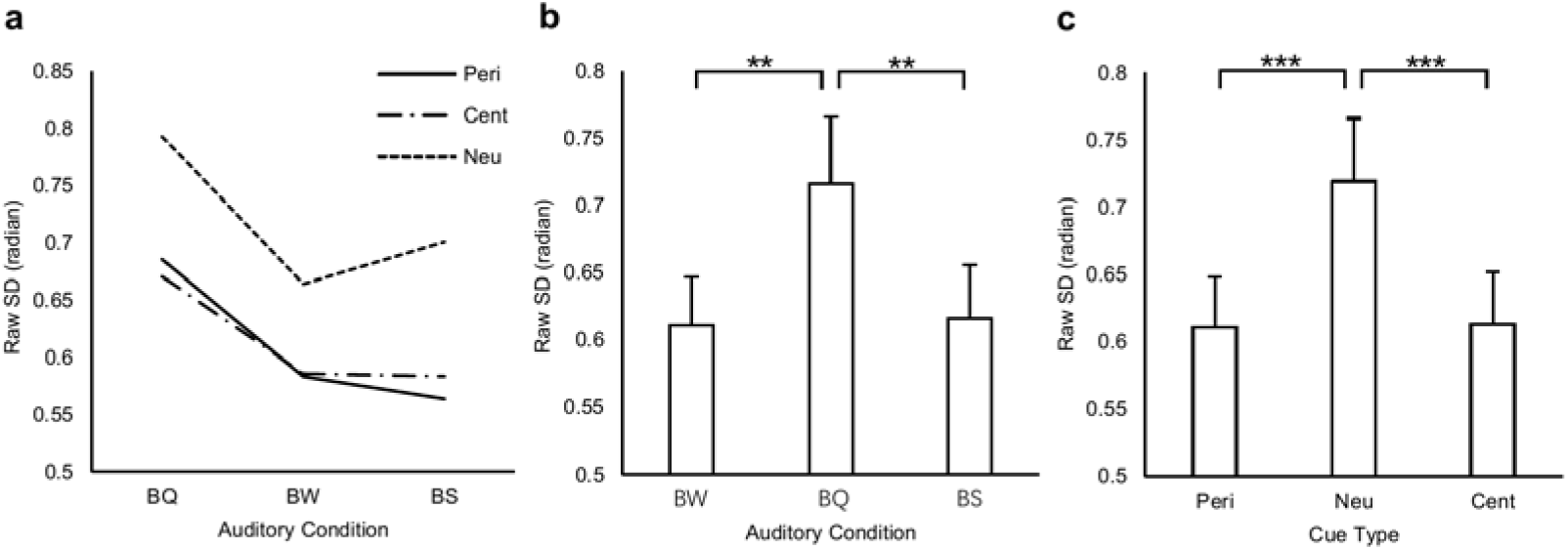
Raw SD results. (a) mean values for each condition. (b) the main effect of auditory condition. (c) the main effect of cue type. Error bars denotes standard errors. Peri = peripheral cue, Neu = no cue, Cent = central cue. \**p* <0.05, \*\**p*< 0.01, \*\*\**p*< 0.001, the same below.

#### Model fitting results

For the guess rate (see Fig. 4a), the main effect of cue type (*F* (2, 38) =1.069, *p*=0.354, η^2^=0.053; Fig. 4c) and the interaction between auditory condition and cue type (*F* (4, 76) =0.146, *p*=0.964, η^2^=0.008) was not significant. However, there was a significant main effect of auditory condition (*F* (2, 38) =3.822, *p*=0.031, η^2^=0.167). The post hoc *t*-tests (see Fig. 4b) showed that only background white noise decreased the guess rate compared to background quiet condition (*t*=-2.602, *p*=0.018), no other difference was found (*ps*>0.05).

**Fig. 4.**
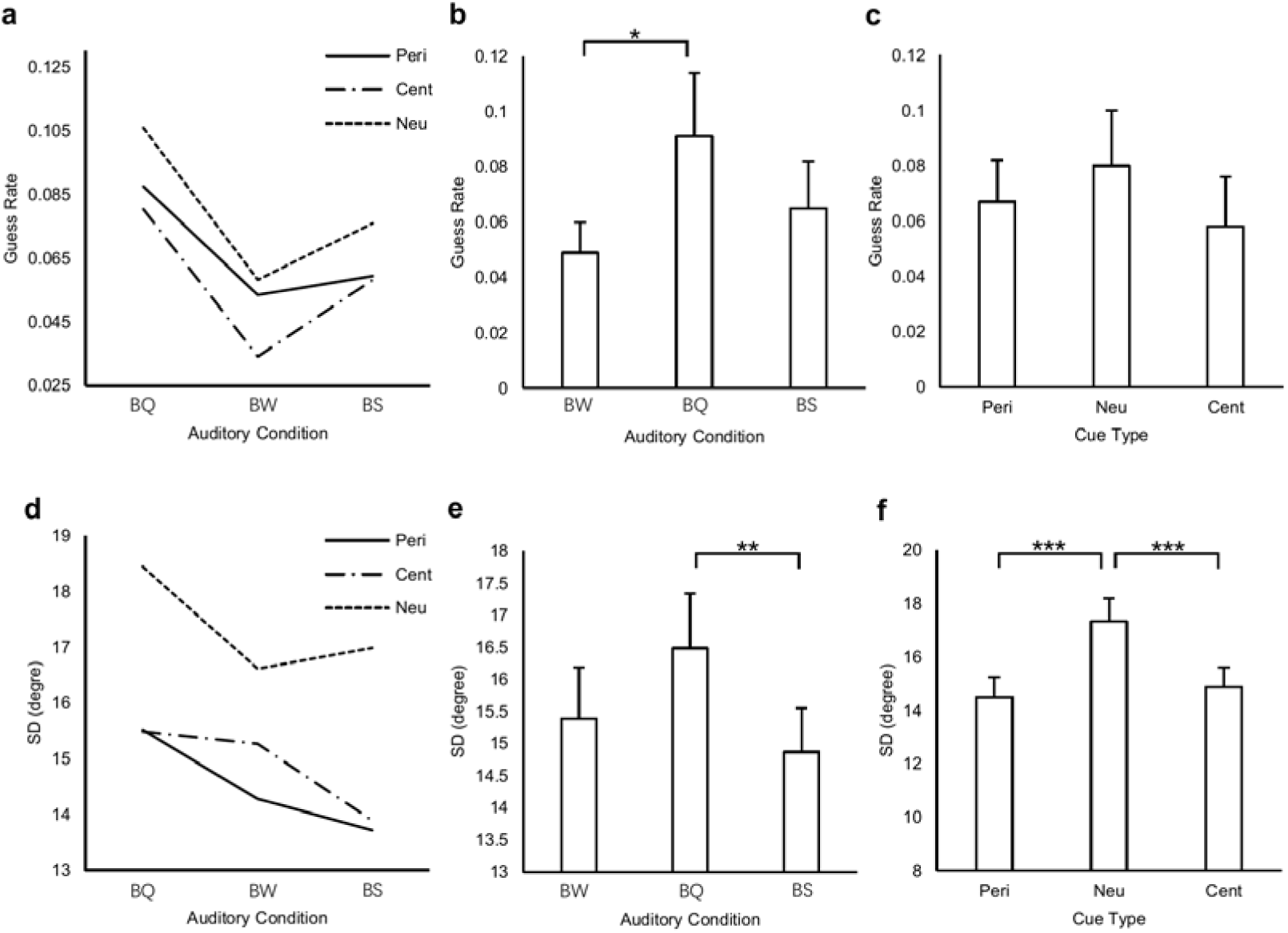
Model fitting results. (a-c) the averaged guess rate for each condition (a), and for the main effect of auditory condition (b) and of cue type (c). (d-f) the averaged SD for each condition (d) and for the main effect of auditory condition (e) and of cue type (f).

For SD (see Fig. 4d), the interaction between auditory condition and cue type was not significant (*F* (4, 76) =0.673, *p*=0.613, η^2^=0.034), but the main effect of auditory condition was significant (*F* (2, 38) =4.299, *p*=0.021, η^2^ =0.185). Post hoc *t*-tests (see Fig. 4e) showed that only background speech noise decreased SD compared to background quiet condition (*t*=-3.753, *p*=0.001), and no other difference was found (*ps*>0.05). In addition, the main effect of cue type was also significant (*F* (2, 38) =15.413, *p*<0.001, η^2^=0.448). Post hoc *t*-tests (see Fig. 4f) showed both peripheral (*t*=-4.919, p<0.001) and central (*t*=-4.590, p<0.001) retro-cues decreased SD compared to no cue condition, there was no significant difference between the two cues (p=0.503).

### Physiological results

for EMG signals, the main effect of auditory condition was not significant (*F* (2, 38) =0.448, *p*=0.642, η^2^=0.023). The background white noise has the lowest EMG values (*M*=-3.679, *SD*=1.305), followed by the background speech noise (*M*=-3.654, *SD*=1.286). The background quiet condition has the highest EMG values (*M*=-3.613, *SD*=1.424).

for EDA signals, the main effect of auditory condition was not significant either (*F* (2, 38) =0.183, *p*=0.834, η^2^=0.01). The background white noise has the highest EDA values (*M*=0.002305, *SD*=0.027888), followed by the background speech noise (*M*=0.002302, *SD*=0.027936). The background quiet condition has the lowest EDA values (*M*=0.002279, *SD*=0.027894).

### The correlation between the behavioral and physiological results

As shown in Fig. 5, the decrease of guess rate induced by background white noise was significantly correlated with the decrease of EMG values (*r*=0.443, *p*=0.05, see Fig. 5a). In addition, the decrease of SD induced by background speech noise was significantly correlated with the increase of EDA values (*r*=-0.489, *p*=0.03, see Fig. 5b). These findings altogether suggested that both background white noise and speech noise led to improvement through arousal enhancement (see supplementary Table 1 for more details).

**Fig. 5.**
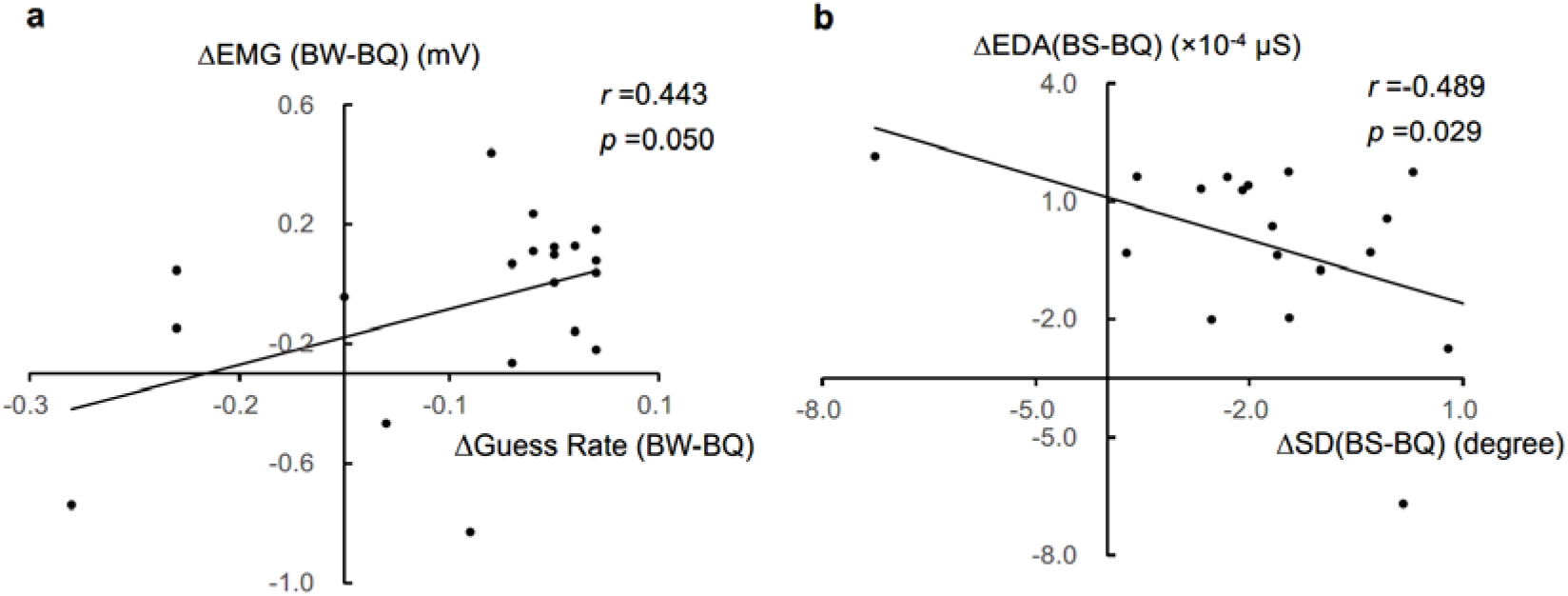
Correlations between behavioral and physiological results. (a) the correlation between the guess rate of background white noise relative to background quiet condition and their EDA difference. (b) the correlation between the SD of background speech noise relative to background quiet condition and their EMG difference.

## Discussion

In the present study, we investigated whether background sound influenced visual WM performance. Our findings suggested that both background speech and white noise facilitated WM performance. Modeling results further revealed that compared with background quiet condition, background white noise lowered the guess rate and background speech facilitated the precision of visual WM.

Physiological recordings indicated that changes in arousal levels might serve as the mechanism underlying these effects. Although we did not observe significant physiological different changes induced by the two kinds of background sound (yet noticing the mean difference), there existed individual differences in behavior that could be accounted by the physiological pattern, which was associated with arousal status. Specifically, the changes in EDA were correlated with the precision difference when background speech was compared with quite; the changes in EMG were correlated with the guess rate difference when background white noise was compared with quiet condition. Taken together, these findings suggested that the beneficial effects of background sound on visual WM was related to the arousal enhancement. However, we here could not exclude an alternative explanation that background sound may affect visual WM performance by interacting with the phonological loop (Baddeley, 1996), although such interaction has been minimized using Gabor patches as stimuli.

In the present study, we also questioned whether background sound would modulate the retro-cueing effects. The answer seemed to be no, since we did not observe an interaction between the types of background sound and types of retro-cues. Then, how to interpret this result? As discussed earlier, the background speech in this experiment did not cause excessive levels of arousal. Instead, it led to relatively low levels of arousal similar to those in white noise conditions. Therefore, participants could benefit from both background speech and white noise by excluding task-irrelevant information. Compared to these two kinds of background sound, spatial retro-cues directly provided task-relevant information and narrowed the attention to a more specific location. In other words, attentional selection might approach a ceiling effect in the presence of spatial cues, which left few spaces for background sound to exclude task-irrelevant information a step further. Another possibility was that the task was in the visual domain and participants could ignore the background sounds from the task irrelevant domain. In this way the task-relevant information selected by spatial retro-cues was not affected by the background sounds.

It should be noted that background speech is often found to impair the cognitive task at hand (Colle & Welsh, 1976; Hellbrueck, Kuwano, & Namba, 1996; Klatte, Meis, Sukowski, & Schick, 2007; Salamé & Baddeley, 1987), but the current study revealed an opposite effect. Another study using an N-back task also found that low-arousal speech improved performance (Han, Liu, Zhang, Jin, & Luo, 2013), which was consistent to our findings. The discrepancy in different tasks indicated that the effects of background sound in environment should not be based on the assumption of equivalency across all situations. Researchers should consider the type of task being performed as well as the arousal level induced by the sound when evaluating its effect.

Our main purpose was to compare different types of background sound, so the sound volume was not varied and set to 55dB. The complexity of background sound could lead to different effects (Baker & Holding, 1993). Moreover, the effects of volume depended on personality of introverts or extraverts (Furnham & Bradley, 1997). Therefore, future study could extend our setting to investigate the effects caused by different sound volumes to individuals with different personalities.

Some studies applied background white noise to enhance cognitive performance in children with attention deficits (for example, ADHD) and proposed a stochastic resonance theory by alter the “signal to noise” ratio through adding different level of background white noise (Helps et al., 2014; Sderlund & Sikstrm, 2012; G. B. Söderlund, Sikström, Loftesnes, & Sonuga-Barke, 2010). It would be interesting to further compare between background speech and white noise on visual WM in special groups to extend our findings out of normal participants.

In addition to background speech, music is another common form of background sound in natural settings. It has been found that listening to music could increase drivers’ arousal and led to improvement in monotonous driving tasks (Ünal, de Waard, Epstude, & Steg, 2013). Future studies could further investigate whether a similar benefit would be found when performing visual WM tasks.

In conclusion, background speech and white noise improved visual WM performance. Specifically, background speech increased the precision of visual WM and background noise lowered the guess rate. Both of them correlated with changes in physiological measurement linked with arousal. Our results suggest that arousal played an important role in explaining the facilitating effects of background sound on visual working memory.

## Acknowledgement

This work was supported by the National Social Science Foundation of China (17ZDA323), the Shanghai Committee of Science and Technology (19ZR1416700), and the Hundred Top Talents Program from Sun Yat-sen University.

## Supplementary

**Table 1.** The correlation between the behavioral and electrophysiological results.

| | | ΔEMG | ΔEDA |
| --- | --- | --- | --- |
| **BW - BQ** | ΔRaw SD | 0.242 | -0.481* |
| | ΔGuess Rate | 0.443* | -0.300 |
| | ΔSD | -0.162 | -0.364 |
| **BS – BQ** | ΔRaw SD | 0.349 | -0.253 |
| | ΔGuess Rate | 0.100 | 0.061 |
| | ΔSD | 0.428 | -0.489* |

*Note.* \* $p < 0.05$ . BW = Background White Noise, BQ = Background Quiet, BS = Background Speech.

